# Viroporin activity from rotavirus nonstructural protein 4 induces intercellular calcium waves that contribute to pathogenesis

**DOI:** 10.1101/2024.05.07.592929

**Authors:** J. Thomas Gebert, Francesca J. Scribano, Kristen A. Engevik, Asha A. Philip, Takahiro Kawagishi, Harry B. Greenberg, John T. Patton, Joseph M. Hyser

## Abstract

Acute gastroenteritis remains the second leading cause of death among children under the age of 5 worldwide. While enteric viruses are the most common etiology, the drivers of their virulence remain incompletely understood. We recently found that cells infected with rotavirus, the most prevalent enteric virus in infants and young children, initiate hundreds of intercellular calcium waves that enhance both fluid secretion and viral spread. Understanding how rotavirus triggers intercellular calcium waves may allow us to design safer, more effective vaccines and therapeutics, but we still lack a mechanistic understanding of this process. In this study, we used existing virulent and attenuated rotavirus strains, as well as reverse engineered recombinants, to investigate the role of rotavirus nonstructural protein 4 (NSP4) in intercellular calcium wave induction using *in vitro*, organoid, and *in vivo* model systems. We found that the capacity to induce purinergic intercellular calcium waves (ICWs) segregated with NSP4 in both simian and murine-like rotavirus backgrounds, and NSP4 expression alone was sufficient to induce ICWs. NSP4’s ability to function as a viroporin, which conducts calcium out of the endoplasmic reticulum, was necessary for ICW induction. Furthermore, viroporin activity and the resulting ICWs drove transcriptional changes indicative of innate immune activation, which were lost upon attenuation of viroporin function. Multiple aspects of RV disease severity *in vivo* correlated with the generation of ICWs, identifying a critical link between viroporin function, intercellular calcium waves, and enteric viral virulence.

## INTRODUCTION

Acute gastroenteritis (AGE) remains a leading cause of mortality and morbidity among young children worldwide^1^. Viruses are the most common enteric pathogens, with rotavirus alone accounting for one quarter of all cases of severe pediatric AGE^2,3^. Viral AGE typically presents with watery diarrhea, vomiting, fever, abdominal pain, and malaise^4^. While most cases of viral AGE are self-limiting, AGE can progress to cause life-threatening dehydration and volume depletion. For such cases, oral rehydration therapy (ORT) is the standard of care^5^. ORT has tremendously reduced the mortality of AGE and all-cause childhood mortality as a result. Yet still today nearly 500,000 children die from AGE annually^2^. While effective to combat dehydration, ORT offers symptomatic treatment without addressing the underlying infection. As such, ORT does not prevent viral spread or outbreaks.

In addition to ORT, the development of live-attenuated rotavirus vaccines has helped to reduce the burden of AGE worldwide. Rotavirus (RV) has long been the most common cause of AGE in children. Both clinical trials and real-world efficacy studies suggest that in the United States, RV vaccines are around 85% effective in the prevention of RV-associated hospitalizations^6,7^. Unfortunately, this degree of efficacy has not been reproduced in areas of the world with higher rates of all-cause childhood mortality. While RV vaccine efficacy is about 86% in countries with low childhood mortality, it falls to just 54-63% in countries with moderate and high rates of childhood mortality^8^. Improved vaccines will be integral to combat RV globally and achieving this goal will require a better understanding of the molecular mechanisms of rotavirus pathogenicity. Thus far, all licensed RV vaccines have been based on live-attenuated strains^9^. While multiple groups are working to develop subunit, mRNA, and inactivated RV vaccines, these efforts have yet to show adequate efficacy in clinical trials. A recently developed subunit vaccine based on the RV spike protein, VP8, failed to meet efficacy criteria in initial stages of clinical trials, and further development was halted^10^. While other candidates have shown promise in pre-clinical studies, live-attenuated vaccines (LAVs) remain the most effective tool for conferring protection against RV.

Despite the widespread use of LAVs, we have yet to fully comprehend the molecular mechanisms that allow for rotavirus attenuation. As such, the development of LAVs for RV has relied on empirical assessment of cell-culture adapted or naturally circulating attenuated strains^11,12^. The advent of the reverse genetics system for RV offers a promising alternative for vaccine development^13^. Currently, simian, some human, and murine-like recombinant RV strains show efficient rescue with the existing reverse genetics strategies, and there have been recent advances toward a higher efficiency human RV reverse genetics system^14,15^. As such, there is potential to intelligently engineer an improved live-attenuated strain. This could allow for attenuation of virulence without major changes to viral antigenicity or fitness, ultimately improving immunogenicity and protection. However, without a comprehensive understanding of rotavirus virulence factors, such rational design is unlikely to generate the desired attenuated traits.

The virulence of RV disease is multigenic, having been directly associated with 5 of the 11 rotavirus genes. These include the genes for the capping enzyme and interferon antagonist, VP3, the outer capsid proteins VP4 and VP7, the interferon-antagonizing protein NSP1, and the multifunctional NSP4 ^16,17^. Broadly, RV virulence is proportional to the severity of diarrhea it causes. RV-induced diarrhea is multifactorial, driven both by malabsorption and hypersecretion^18^. Our understanding of RV disease evolved upon the discovery of the enterotoxin function of NSP4^19^. In a 1996 study, investigators found that intraperitoneal injection of purified NSP4 was sufficient to induce diarrhea in mouse pups^19^. This study established NSP4 as the first known enterotoxin of viral origin. However, studying the potential enterotoxin activity of NSP4 during a *bona fide* RV infection has proven more challenging. While the enterotoxin NSP4 is likely a key mediator of RV disease, we have yet to determine its relative contribution to the overall diarrhea phenotype. NSP4 has since been found to also function intracellularly as a viroporin, forming a calcium (Ca^2+^)-conductive ion channel that releases Ca^2+^ from the endoplasmic reticulum (ER) store^20,21^. NSP4 variants associated with attenuated diarrhea induction *in vivo* show a diminished capacity to increase cytosolic Ca^2+^ when expressed in Sf9 cells, suggesting a role for NSP4 Ca^2+^ mobilization in virulence^22^.

We recently discovered that RV-infected cells release adenosine diphosphate (ADP) in pulses throughout infection, constituting a previously unidentified contributor to RV virulence^23^. By activating the P2Y1 receptor, extracellular ADP drives an increase in cytosolic Ca^2+^ in neighboring, uninfected cells, producing a signal known as an intercellular Ca^2+^ wave (ICW). Blocking the P2Y1 receptor, and the resulting ICWs, reduces disease severity in neonatal mouse pups infected with a heterologous (simian) RV strain. Furthermore, preventing ICWs reduces viral spread in cell culture^24^. Importantly, knockdown of NSP4 reduces the frequency of ICWs from RV-infected cells, suggesting that a function of NSP4 may be involved in ICW induction.

Given these findings, it is likely that ICWs contribute to RV virulence; however, we have yet to fully characterize the viral determinants of this signaling. In this study, we used existing human and porcine RV strains, as well as novel recombinants generated by the RV reverse genetics system, to examine the role of NSP4 in the induction of ICWs using *in vitro*, organoid, and *in vivo* model systems. Ultimately, we found that the ability of RV to generate ICWs segregates with NSP4, expression of NSP4 alone is sufficient to generate ICWs, and multiple aspects of RV disease severity correlate with the ability to generate ICWs. Overall, this work implicates NSP4 viroporin activity as the primary driver of RV-induced ICWs and a critical component of the pathogen-induced changes in host cell physiology.

## RESULTS

### Virulent porcine RV induces a higher frequency of ICWs than its attenuated counterpart

Based on our previous studies which demonstrated that pharmacologic blockade of ADP-mediated ICWs lessened disease severity in RV-infected mice^23^, we sought to determine whether attenuated RV strains elicit fewer ICWs during infection. We performed long-term Ca^2+^ imaging on monolayers infected with RV strains of varying virulence. For these studies, we first investigated the porcine rotavirus strains OSUv and OSUa^25^. OSUv was isolated from a sick neonatal piglet and was serial passaged in cell culture to generate the attenuated strain, OSUa ^22,25–27^. Upon imaging, we found that cells infected with OSUv induced more ICWs than OSUa-infected cells (Figure 1A-D). This was quantifiable by two distinct methods. First, we infected monolayers of MA104 cells expressing GCaMP6s (MA104-GCaMP6s) with the indicated RV strains at MOI 0.01. For each infection, we imaged a 10x10 mm stitched field-of-view once every 2 min for 30 min beginning at 18 hpi. We then generated maximum intensity projections of each field-of-view, allowing for visualization of all ICWs detected in the 30 min period (Figure 1a). Additionally, to capture a longer duration of the infection, we infected MA104-GCaMP6s monolayers with the indicated RV strains at MOI 0.01 and imaged 8 positions per monolayer every 1 minute from 8-24 hpi. We developed an analysis pipeline in FIJI that allowed for automated quantitation of ICWs detected within each field-of-view during the 16 hrs of imaging. Using this approach, we detected a 4-fold increase in ICWs in monolayers infected with OSUv compared to those infected with either mock inoculum or OSUa (Figure 1bi). Furthermore, the ICWs from OSUv-infected cells began earlier after infection than those from OSUa-infected cells (Figure 1bii). The overall increase in Ca^2+^signaling associated with OSUv is further illustrated by Ca^2+^ traces from a representative mock-, OSUv-, and OSUa-infected cells (Figure 1c).

**Figure 1:**
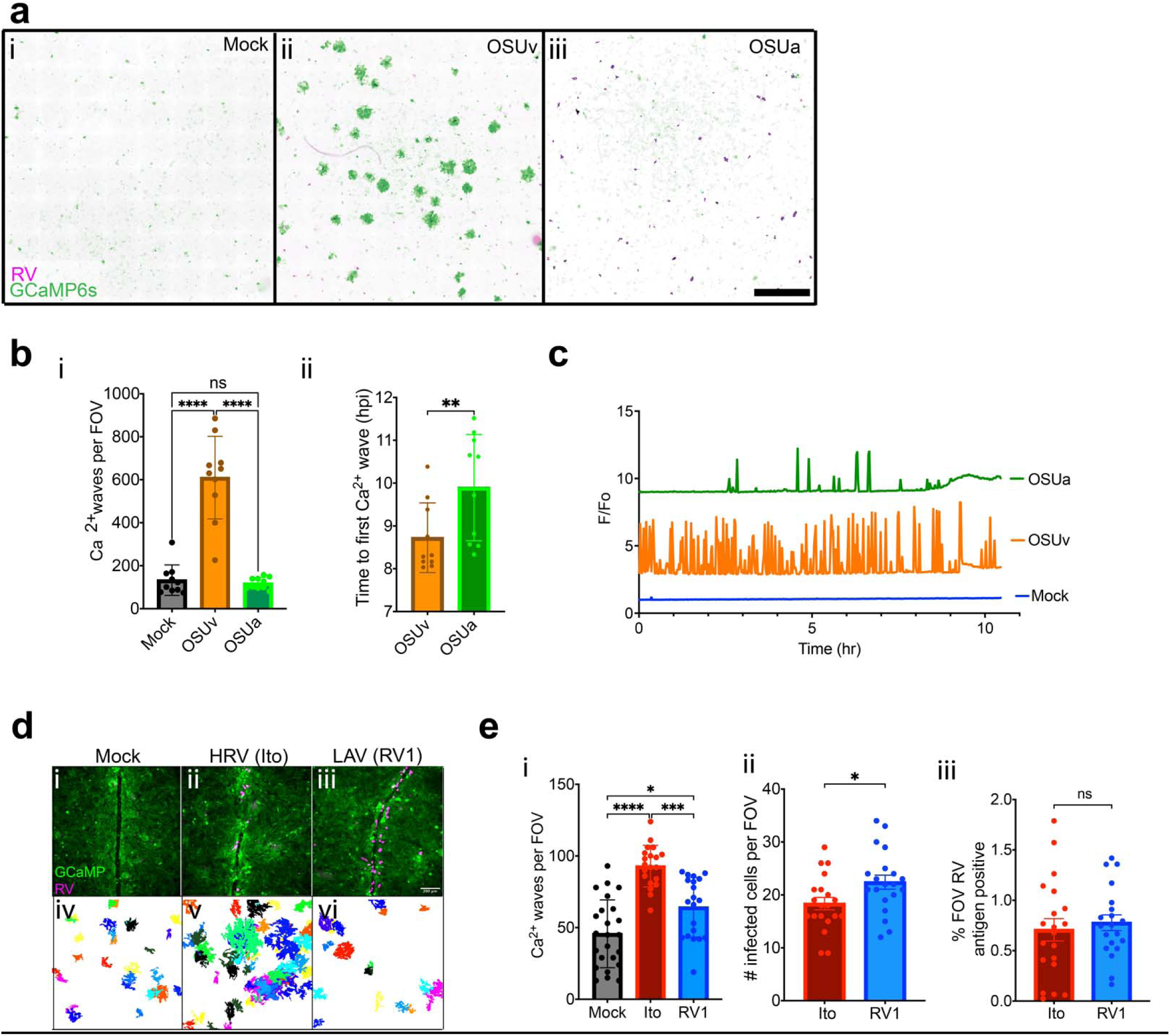
Frequency of intercellular Ca^2+^ waves (ICWs) is associated with virulence. **a**) Maximum intensity projections of MA104-GCaMP6s monolayers imaged after infecting with mock (**i**), OSUv (**ii**), or OSUa (**iii**) rotavirus at MOI 0.01. ICWs are indicated by regions of high GCaMP6s intensity (green) surrounding RV-infected cells (magenta), seen predominantly in the OSUv-infected monolayer. 10x10 mm regions of each monolayer were imaged for 30 min beginning 18 hrs post-infection. **b**) MA104-GCaMP6s monolayers were infected at MOI 0.005 and imaged from 8-24 hpi. ICWs were quantitated using an automated pipeline developed in ImageJ. The quantity (**i**) and latency (**ii**) of ICWs between mock, OSUv, and OSUa infection. Representative data from one of three biological replicates. **c**) representative intracellular Ca^2+^ traces from mock-, OSUv-, or OSUa-infected cells. **d**) Representative human intestinal organoid monolayers expressing GCaMP6s (green) and infected with mock inoculum **(i),** virulent human RV (**ii**), or an attenuated RV strain (**iii**). Monolayers were imaged from 8-24 hpi before fixing and staining for RV antigen (magenta). ICWs detected over the course of imaging were arbitrarily pseudo-colored and projected into a single image for each multipoint (iv-vi). e) The quantity of ICWs (**i**) and infected cells (**ii**), and the percentage of monolayer area positive for RV antigen compared between mock, Ito, and RV1 infection. Data combined from 3 biological replicates. Data in bar charts represent mean with error bars denoting SD. *p<0.05, **p<0.01, ***p<0.001, ****p<0.0001.

### Virulent human RV strain induces higher frequency of ICWs than live-attenuated strain

Given the association between virulence and ICWs observed with the porcine RV strains, we next aimed to determine whether this relationship would hold among human RV strains. Using a similar experimental approach, we infected monolayers of jejunal human intestinal organoids (jHIOs) engineered to express the cytosolic calcium indicator GCaMP6s (jHIO-GCaMP6s) with virulent or attenuated human RV strains. For the virulent strain we selected *Ito*, as it was previously shown to infect HIEs and induce ICWs^23,28^. For the attenuated strain we selected RV1, which is one of the most widely used live-attenuated vaccines (Rotarix)^9^. Representative fields-of-view for mock-, Ito-, and RV1-infected jHIO-GCaMP6s monolayers show the expected detection of RV near the site of a monolayer scratch (Figure 1di-iii). The ICW phenotypes associated with each virus are illustrated by the projection of all ICWs detected in each field-of-view over the preceding 18 hrs of imaging, which have been pseudo-colored to distinguish individual ICW events (Figure 1div-vi). Quantification of ICWs detected across all fields-of-view confirmed that, consistent with observations using the porcine strains, the virulent human strain, Ito, induced significantly more ICWs during infection than the attenuated strain, RV1 (Figure 1ei). To determine whether this may have been due to increased infectivity of the virulent strain, we fixed monolayers post-imaging and detected RV antigen by immunofluorescence. We quantitated the number of infected cells per field-of-view (Figure 1e ii) and the percentage of each field-of-view that was antigen positive after flat-field correction, background subtraction, and Otsu thresholding^29^ (Figure 1e iii). The number of infected cells was comparable between groups, with slightly more infected cells in monolayers infected with RV1 (18.4 vs. 22.4 for RV1 and Ito, respectively, Figure 1eii). We detected no difference in the percentage of the monolayer area that was antigen positive (Figure 1eiii). Taken together, the greater number of ICWs in monolayers infected with the virulent human strain, Ito, was not attributable to increased infectivity.

### ICW phenotype segregates with gene 10, encoding nonstructural protein 4 (NSP4)

We next aimed to determine which viral protein(s) may drive the observed difference in ICW phenotype. For these experiments we focused on the OSUv and OSUa, as high sequence homology facilitates phenotype-genotype mapping^22,30^. In previous studies, we found that using a small-hairpin RNA to abrogate the expression of NSP4 reduced the number of ICWs that occur during infection^23^. Based on this, we hypothesized that differences in NSP4 account for the observed differences in the ICW phenotype between OSUv and OSUa. To isolate the effects of the changes in NSP4 between OSUv and OSUa, we generated two novel RV strains using the reverse genetics system, both in an SA11 (simian RV) background. In the first strain, we replaced SA11 gene 10, which encodes NSP4, with that from OSUv (SA11-G10-OSUv). In the second, we replaced gene 10 with that from OSUa (SA11-G10-OSUa) (Figure 2ai). In both strains, we used the previously characterized gene 7 construct encoding RV NSP3 and the fluorescent protein mRuby, which facilitates identification of infected cells during imaging^23^. Performing long-term Ca^2+^ imaging of infected MA104-GCaMP6s monolayers revealed that SA11-G10-OSUv elicited a greater number of ICWs than did SA11-G10-OSUa (Figure 2b), suggesting NSP4 is a critical determinant of ICWs.

**Figure 2:**
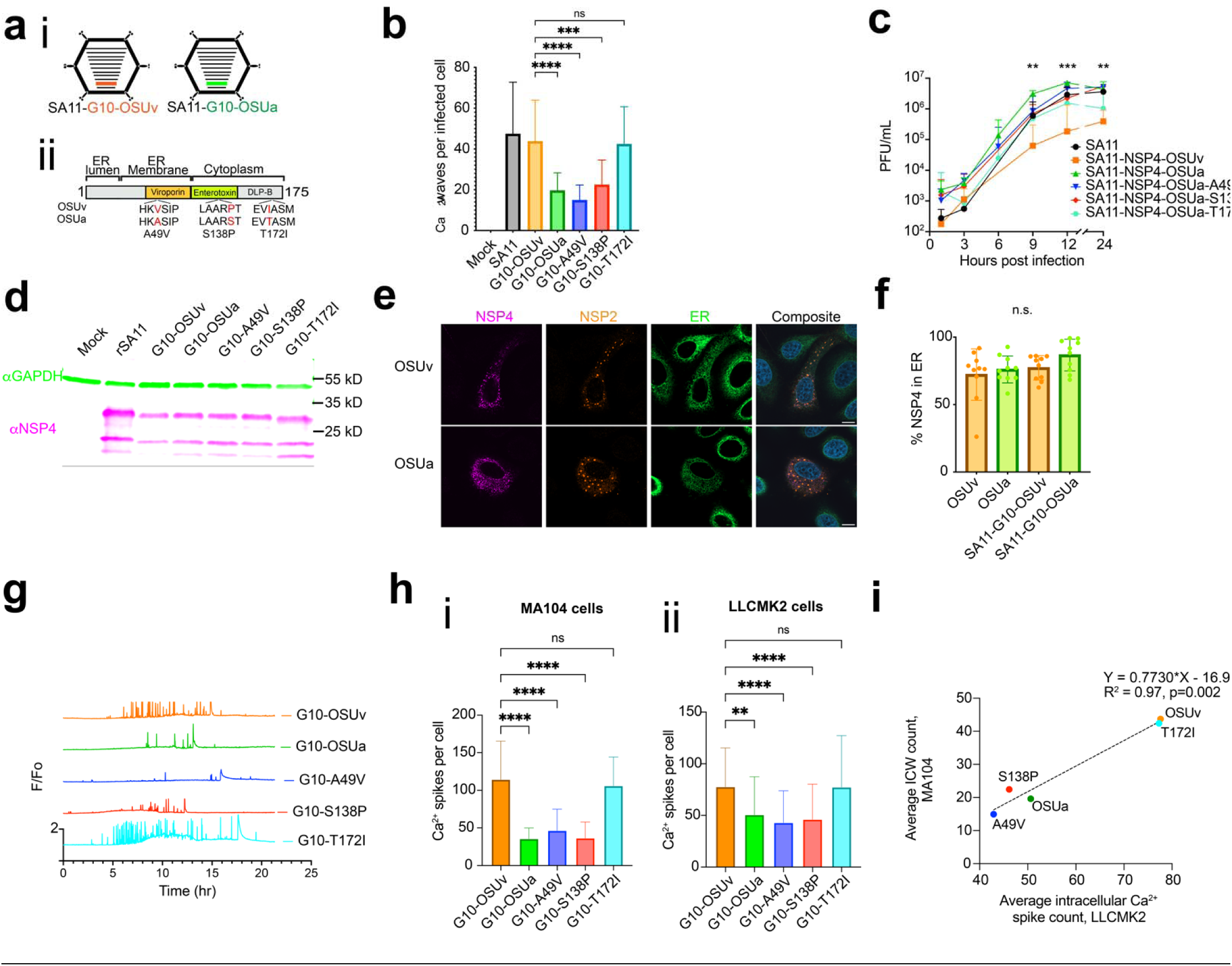
The difference in ICW phenotypes is attributable to NSP4. **a)** Schematic representation of the recombinant SA11 strains with gene segment 10 encoding NSP4 from either OSUv or OSUa (i). The linearized domain map (ii) illustrates the 3 amino acid differences between OSUv and OSUa NSP4. **b**) Quantification of ICWs in MA104-GCaMP6s monolayers infected with mock inoculum, SA11, SA11 expressing gene 10 (NSP4) from OSUv or OSUa, or SA11 expressing the NSP4 from OSUa with one of the 3 amino acids reverted to the OSUv-NSP4 identity in turn. Monolayers were imaged from 8-24 hpi and ICWs quantitated across 3 biological replicates. **c**) Growth curves comparing viral yield at 6 time points post-infection. **d**) NSP4 detected at 9 hpi in MA104 monolayer lysate following infection at MOI 5 with the indicated strain of recombinant SA11. **e**) Representative images of SA11-G10-OSUv and -OSUa infected monolayers expressing ER-localized GFP (green) with NSP4 (magenta) and NSP2 (orange) detected by immunofluorescence at 12 hpi. Scale bar = 10μm. **f**) Mander’s coefficient to estimate NSP4 colocalization with Sec61β as a surrogate for ER localization. Combined data from 15 cells across 3 biological replicates. **g)** Representative intracellular Ca^2+^ traces from cells infected with the indicated strain of recombinant SA11. **h)** Intracellular Ca^2+^ spike counts from (i) MA104-GCaMP6s or (ii) LLCMK2 cells (ICW deficient) infected with the indicated strain of recombinant SA11. Combined data from 3 biological replicates. **i**) Linear regression estimating the association between intracellular Ca^2+^ spike counts in LLCMK2-GCaMP6s cells and ICW counts in MA104 GCaMP6s cells.

### The C-terminal domain of NSP4 is responsible for the differing ICW phenotypes of OSUv and OSUa

Through sequencing we found that NSP4-OSUv and NSP4-OSUa differ at just three amino acids at positions 49, 138, and 172 (Figure 2aii, Supplemental Figure 1). To determine which of these changes was responsible for the differing ICW phenotypes, we generated an additional three strains of recombinant SA11. Each of these encoded NSP4 from OSUa, but with one of the three amino acids reverted to the OSUv sequence in turn. Only the T172I reversion was sufficient to recover the ICW phenotype (Figure 2b), suggesting the C-terminal domain of NSP4 may play a role in activating the release of ADP to drive ICWs. We next aimed to determine whether the C-terminal domain of NSP4 acted directly upstream of ICWs, or whether the change at amino acid 172 indirectly attenuated ICWs. We reasoned that the changes in the C-terminus of NSP4 may *indirectly* influence waves by (1) decreasing viral replication, (2) decreasing the expression of NSP4, (3) altering the expression of other viral proteins^31^, or (4) by changing the trafficking dynamics of NSP4^32^.

To determine whether the strains lacking ICWs had reduced replication kinetics, which may indirectly cause a decrease in ICWs, we generated growth curves for each of the recombinant SA11 strains in MA104 cells. We observed no reduction in viral replication among the non-wave-inducing strains SA11-G10-OSUa, A49V, or S138P when compared to the wave-inducing strains (Figure 2c). In fact, while there was no difference in the viral yield between 1 and 6 hpi, SA11-G10-OSUa yield was higher at 9, 12, and 24 hpi than that of SA11-G10-OSUv. Thus, the reduction in waves was associated with G10-OSUa was not likely a result of an overall decrease in the rate of viral replication. We next examined the relative levels of NSP4 between viral strains. There was no difference in the level of NSP4 detected by western blot between SA11-G10-OSUv and -OSUa, though levels trended higher for SA11-G10-OSUa (Figure 2d, Supplemental Figure 2). As such, the reduction in waves could not be attributed to a reduction in the level of NSP4.

Finally, we aimed to assess whether NSP4-OSUv and NSP4-OSUa differed in cellular localization or trafficking. NSP4 is first synthesized in the ER, where it is co-translationally inserted into the ER membrane. NSP4 conducts ER Ca^2+^ into the cytoplasm, which activates autophagy^20,21,33^. NSP4, in association with Sec24 and LC3, then exits the ER in COPII vesicles which eventually associate with viroplasms^33,34^. We aimed to determine whether ER exit may occur at a different time post-infection between NSP4-OSUv and NSP4-OSUa. For these studies, we developed a line of MA104 cells (MA104-ER-GFP) that constitutively express green fluorescent protein (GFP) anchored to the transmembrane domain of Sec61β, functioning as a fluorescent marker of the endoplasmic reticulum. We infected MA104 cells with OSUV, OSUa, SA11-G10-OSUv, or SA11-G10-OSUa at MOI 0.1 and fixed 12 hours post-infection, as this is around the time that SA11-G10-OSUv begins to robustly activate ICWs. We did not detect a difference in the proportion of NSP4-OSUv and NSP4-OSUa that co-localized with GFP, suggesting no major difference in the amount of NSP4 retained in the ER membrane at 12 hpi (Figure 2 e-f). Similarly, we found no difference in the percent of NSP4 co-localized with immunofluorescently tagged Sec61β, nor with NSP2 as a marker of viroplasms, the target of NSP4 after it leaves the ER (Supplemental Figure 3).

### Ca^2+^ signaling within infected cells is associated with the ICW phenotype

Previous studies have implicated *intra*cellular Ca^2+^ dysregulation in the initiation of ICWs^35^. We sought to determine whether dysregulation of Ca^2+^ signaling *within* the infected cell may differ between cells infected with a wave-inducing versus a non-wave-inducing strain. The cytosolic Ca^2+^ traces from cells infected with the wave-inducing SA11-G10-OSUv and SA11-G10-T172I seemed to show more dramatic dysregulation relative to strains associated with fewer ICWs (Figure 2g). Indeed, the strains associated with fewer ICWs also show fewer intracellular Ca^2+^ spikes (Figure 2hi). However, ADP signaling is expected to have both a paracrine and autocrine effect. Thus, a virus that induces fewer ICWs would be expected to induce fewer Ca^2+^spikes in the infected cell due to reduced autocrine signaling. To determine whether the strains that induce fewer ICWs may be independently inducing fewer intracellular Ca^2+^ spikes, we used the LLC-MK2 line of monkey kidney epithelial cells. LLC-MK2 cells do not express the P2Y1 receptor and therefore do not show ICWs during RV infection^24^. The difference in intracellular Ca^2+^ spikes within infected cells persists in this cell line, suggesting a P2Y1-independent difference in intracellular Ca^2+^ signaling (Figure 2Hii). Furthermore, across the RV strains, the average number of Ca^2+^ spikes within infected LLC-MK2 cells exhibited a positive correlation with the average number of ICWs in MA104 cells (R^2^=0.97, p=0.002, Figure 2i).

### NSP4 is sufficient to induce ICWs

Given the observed segregation of the ICW phenotype with NSP4, we aimed to determine whether NSP4 would be sufficient to induce ICWs outside of the context of a RV infection. To recapitulate the high levels of NSP4 expression characteristic of RV infection, we turned to adeno-associated virus (AAV) vectors. We generated constructs encoding SA11 NSP4 along with a fluorescent reporter (AAV2-SA11-NSP4-mScarlet, Figure 3a) and transduced MA104-GCaMP6s cells before performing long-term Ca^2+^ imaging. Detection of mScarlet fluorescence confirmed transduction (Figure 3b, top row, magenta). Segmentation and projection of ICWs detected in 30 min for each stitched field-of-view illustrated an upregulation of ICWs associated with SA11-NSP4 (Figure 3b, bottom row). ICW quantitation revealed that SA11-NSP4 transduction elicited a greater number of ICWs relative to AAV2-mScarlet alone or non-transduced controls (Figure 3c). We also generated constructs encoding NSP4 from both OSUv and OSUa. Similar to our findings from our recombinant RV strains, OSUv NSP4 induced a greater number of ICWs than OSUa NSP4 (Figure 3c). Likewise, intracellular Ca^2+^ traces from representative cells suggest that AAV-expressed OSUa NSP4 induces fewer intracellular Ca^2+^ signals than OSUv NSP4 (Figure 3d). Western blot confirmed the expression of NSP4 from the AAV constructs (Figure 3e).

**Figure 3.**
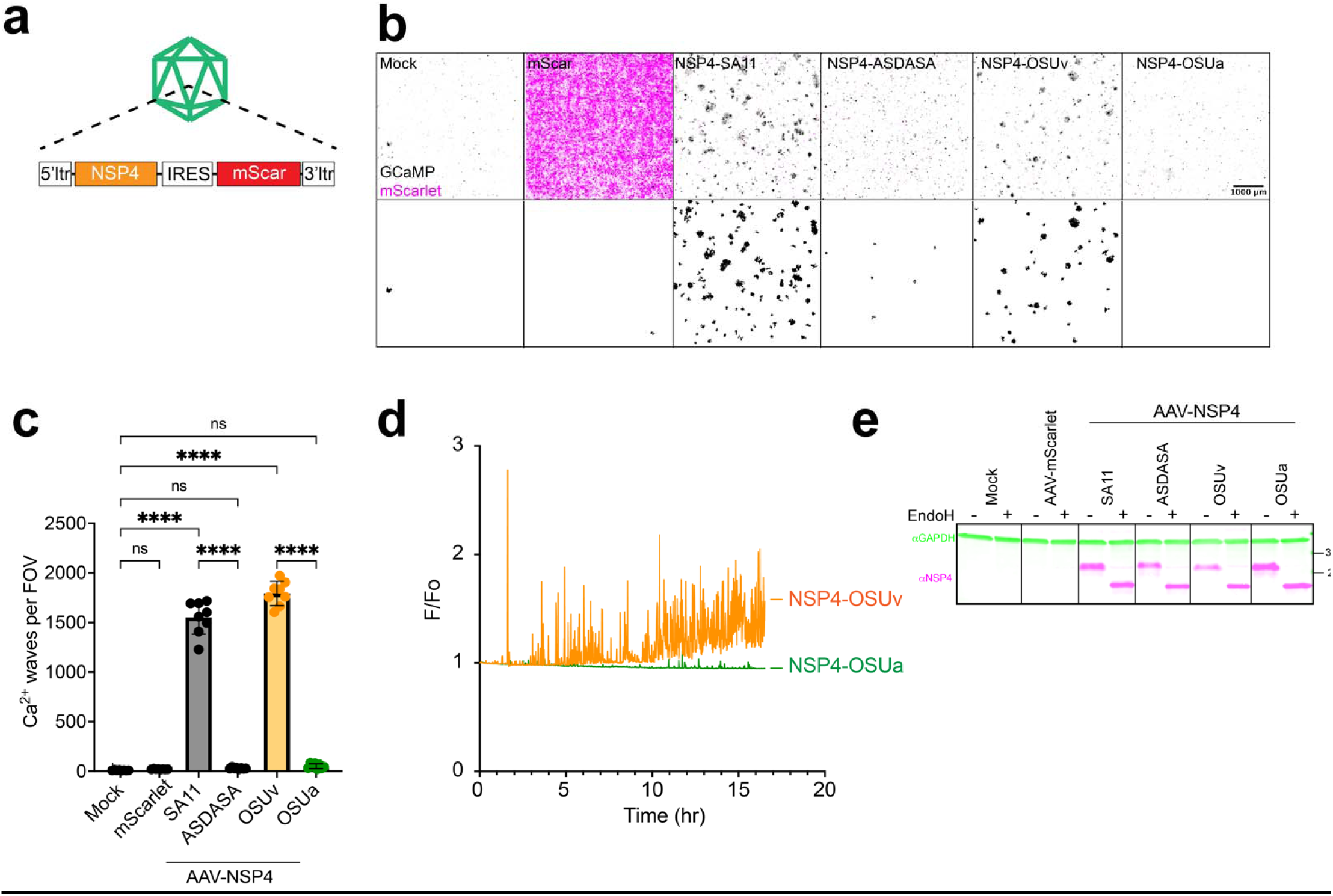
NSP4 is sufficient for ICWs. **a**) Schematic representation of adeno-associated viral vectors used to transduce NSP4 and the fluorescent protein, mScarlet. **b**) mScarlet signal (magenta) overlaid on maximum intensity projections of GCaMP6s signal (black) during 30 min of imaging beginning 48 hours post-transduction (top row). The ICWs post-segmentation (bottom row). **c)** ICWs detected during 18 hrs of imaging following transduction with the indicated AAV constructs. (mScarlet, magenta). Monolayers were imaged every 1 min for 30 min, beginning 52 hrs post transduction. Representative data from one of three biological replicates. **d**) Representative intracellular Ca^2+^ traces from cells expressing OSUv (orange) or OSUa (green) NSP4. **e**) NSP4 detected by western blot in MA104 cells transduced with the indicated AAVs.

### Intact viroporin activity is necessary for NSP4-induced ICWs

Expression of NSP4 by AAV2 transduction offered a flexible platform to explore the mechanistic link between NSP4 and ICWs. NSP4 is a known viroporin that conducts Ca^2+^ out of the ER in infected cells^21^. Given the noted correlation between the frequency of Ca^2+^ signals within infected cells and the frequency of ICWs, we hypothesized that the reduced ICW count associated with NSP4-OSUa may be due to a reduced capacity to disrupt cytosolic Ca^2+^ within the infected cell. To further test this hypothesis, we generated an AAV construct encoding a mutant NSP4 that lacks viroporin activity. This mutant, dubbed the “NSP4-ASDASA” mutant, harbors missense mutations that alter six key residues within the predicted amphipathic domain (IFNTLL ➔ ASDASA)^20^. We have attempted to rescue RV strains with the ASDASA NSP4 mutant via reverse genetics but have never been successful. This is consistent with previous observations suggesting that NSP4-mediated Ca^2+^ dysregulation is critical for RV replication^33^. The AAV platform circumvents this limitation, allowing high expression of NSP4 without viral replication.

Transduction of MA104-GCaMP6s cells with an AAV encoding NSP4-ASDASA and a fluorescent reporter showed no difference in ICWs relative to those transduced with the fluorescent reporter alone (Figure 3c). Western blot with EndoH treatment confirmed that the ASDASA mutant is both expressed and glycosylated, consistent with wildtype SA11 NSP4 (Figure 3e). This finding supports the hypothesis that viroporin activity is necessary for NSP4s ability to induce ICWs. However, quantitation by western blot suggested that the amount of the NSP4-ASDASA may have been slightly lower than that of SA11-NSP4 (Figure 3e). To determine whether the lack of ICW induction by NSP4-ASDASA may have been attributable to decreased protein level, we repeated the imaging experiment using lower doses of AAV-SA11-NSP4 to normalize the protein level to that of NSP4-ASDASA. Even with 10-fold fewer genome copies, SA11-NSP4 maintained the ability to induce ICWs (Supplemental Figure 4a). At this multiplicity, we detected less SA11-NSP4 than ASDASA-NSP4 by western (Supplemental Figure 4b), suggesting that the inability of ASDASA-NSP4 to induce ICWs does not stem from a reduction in the amount of protein. Expression of the fluorescent reporter paralleled these findings (Supplemental Figure 4c).

### NSP4 viroporin activity drives transcriptional changes

Previous studies have shown that RV infection induces broad transcriptional changes within the intestinal epithelium^36^. *How* RV instigates these changes remains an outstanding question. To determine whether NSP4 and its resultant Ca^2+^ signaling affect transcriptional regulation, we performed bulk RNA sequencing on monolayers transduced with the AAVs described above. To differentiate the effects of ICWs and intracellular Ca^2+^ signals, we included cells transduced with SA11 NSP4 and treated with the P2Y1 inhibitor, BPTU, which blocks ICWs^23^. Transcriptional changes in BPTU-treated monolayers should therefore represent the effects of NSP4-induced intracellular Ca^2+^ signals rather than those dependent on ICWs.

Quantification of the mScarlet reporter confirmed transduction of all constructs, albeit with varying degrees of efficiency (Figure 4a). Two-component principal component analysis of transcript counts revealed 3 primary clusters: mock-transduced controls, samples transduced with the vehicle alone or with attenuated NSP4, and samples transduced with virulent NSP4 (Figure 4b). Interestingly, OSUa NSP4 clustered with NSP4 ASDASA and the mScarlet (vehicle) control, whereas OSUv NSP4 clustered with SA11 NSP4. The BPTU treated SA11 NSP4 sample showed more similarity to the untreated SA11 or OSUv NSP4 samples than the ASDASA or OSUa NSP4 samples, suggesting that the ICWs are not the major contributor to the observed transcriptional changes. Instead, this suggests that the *intra*cellular Ca^2+^ dysregulation from NSP4 is the critical effector or the transcriptional changes induced by ICWs are largely similar to those caused by the NSP4-induced Ca^2+^ signals.

**Figure 4:**
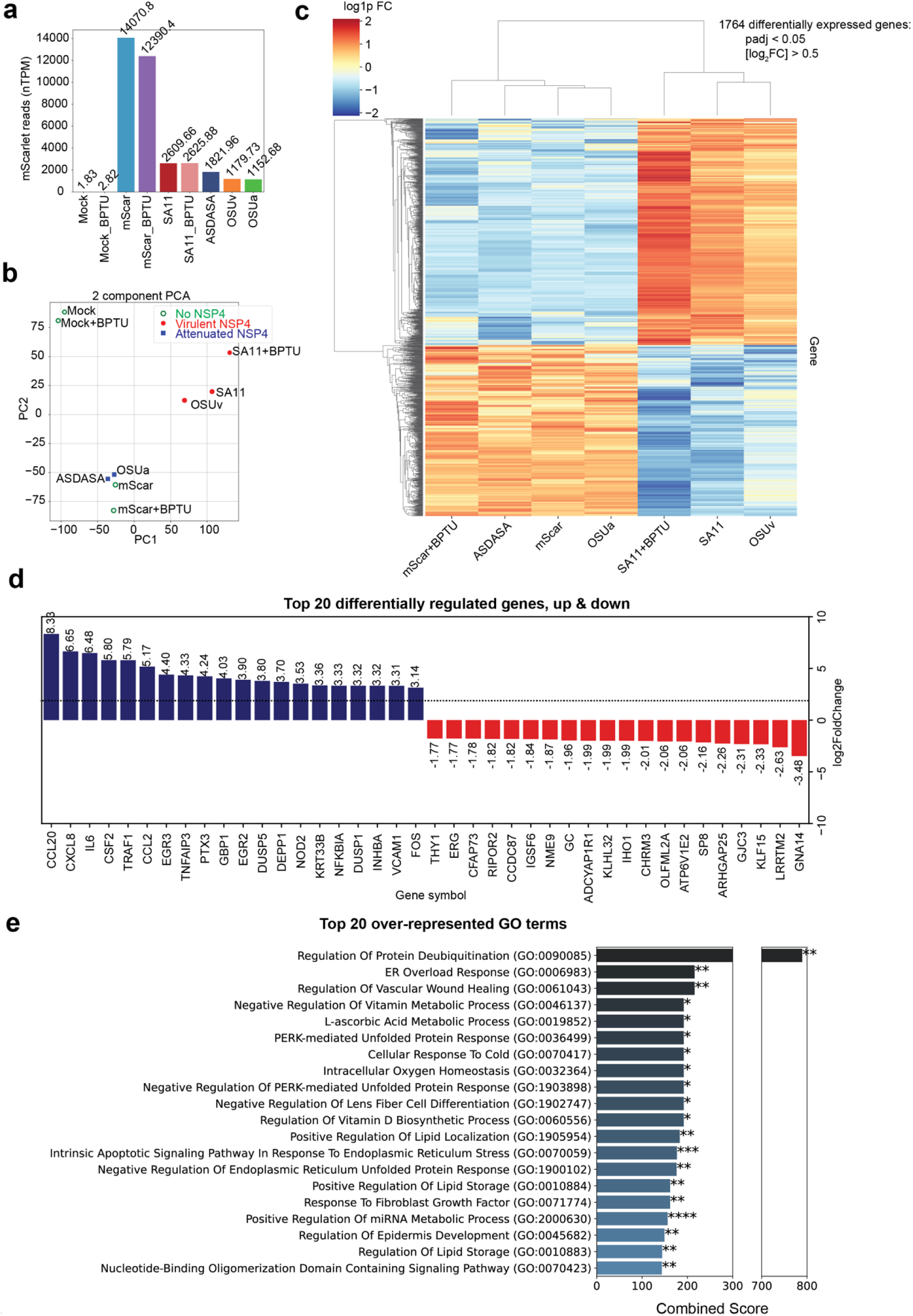
NSP4 viroporin activity drives transcriptional changes. MA104 cells were transduced with 50,000 genome copies of AAV encoding the indicated NSP4 and the fluorescent protein mScarlet. RNA was extracted at 36 hrs post-transduction. **a)** Normalized read counts of mScarlet, as a surrogate of AAV transgene expression. **b**) Two-component principal component analysis plot. **c**) Heat map of differentially regulated genes between ICW-inducing and non-ICW inducing AAV-NSP4 variants and controls. **d**) Top 20 differentially regulated genes. **e**) Overrepresented gene ontology (GO) terms in MA104 cells transduced with ICW-inducing NSP4 variants relative to those transduced with non-ICW inducing NSP4 variants.

Further analysis revealed a total of 1764 differentially expressed genes between samples transduced with no/attenuated NSP4 versus those transduced with virulent NSP4 (padj < 0.05, [log_2_FC]>0.5, Figure 4c), with many of the most highly upregulated genes relating to innate immunity, including CCL20, CXCL8, IL6, CSF2, and TRAF1 (Figure 4d). Importantly, this upregulation was not seen in samples transduced with NSP4 OSUa or the NSP4 ASDASA mutant. Gene set enrichment analysis revealed additional gene ontology terms overrepresented in the samples transduced with virulent viroporins, including genes involved in protein deubiqiutination, ER overload, and PKR-like protein kinase (PERK) regulation (Figure 4e).

### NSP4 drives differences in Ca^2+^ signaling and swelling in RV-infected HIOs

While MA104 cells provided a valuable tool for initial investigation, they are of neither human nor intestinal origin. HIOs provide a RV-susceptible model system that more closely recapitulates the human gastrointestinal epithelium. We first verified that HIOs were susceptible to the recombinant strains of SA11 expressing OSUv or OSUa NSP4 by performing yield assays and immunofluorescence. After a 1-hour inoculation in suspension and 24 hours of growth thereafter, RV antigen was detected by IF in HIOs inoculated with either strain of recombinant SA11 (Figure 5a). Further, RV yield at 24 hpi confirmed replication of both strains in HIOs (Figure 5b).

**Figure 5:**
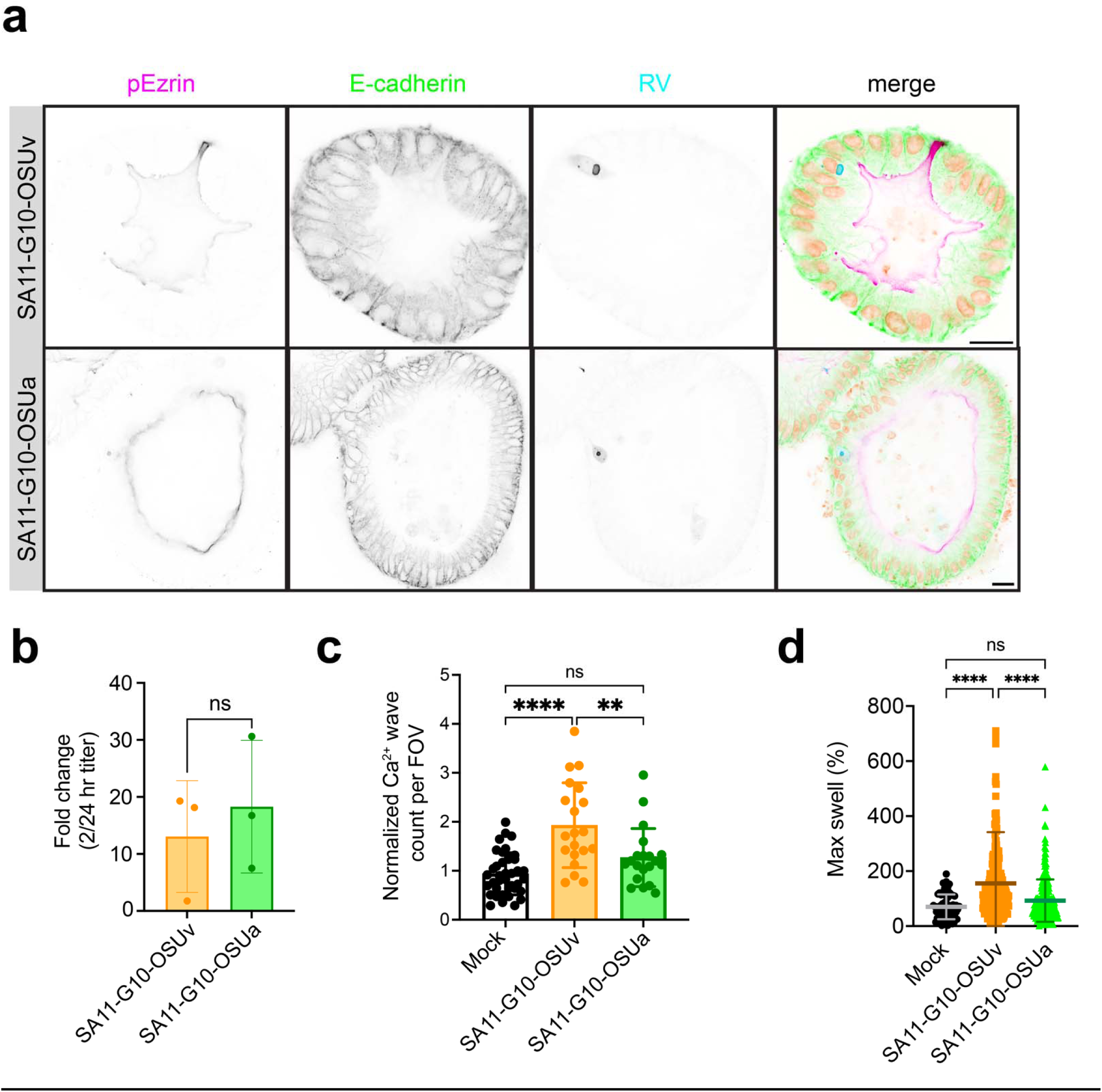
Human intestinal organoids (HIOs) infected with SA11-G10-OSUv and SA11-G10-OSUa support replication and show distinct pathogenic markers. **a**) Representative immunofluorescence images of HIOs infected with SA11-G10-OSUv and -OSUa at 18 hpi. Scale bar = 20μm. **b**) Results from 24 hr yield assay showing comparable yield of SA11-G10-OSUv and -OSUa in HIOs. **c**) ICWs detected in HIO-GCaMP6s monolayers infected with the indicated viruses. Monolayers were imaged every 1 min from 8-24 hpi. **d)** Percent increase between initial and maximum cross-sectional area in 3-dimensional HIOs during infection with mock inoculum, SA11-G10-OSUv or SA11-G10-OSUa. Data combined from 3 biological replicates.

Next, we next aimed to characterize the ICW phenotype associated with each strain in HIOs. For these experiments, we used monolayers generated from the jHIO-GCaMP6s line^37^. We infected differentiated monolayers with RV and performed live Ca^2+^ imaging for 18 hours post-infection. ICW analysis revealed that, similar to our findings in MA104 cells, SA11-G10-OSUv infection induced more ICWs than SA11-G10-OSUa (Figure 5C).

HIOs also provided a valuable model system to investigate fluid secretion, a key aspect of RV pathogenesis. We performed swelling assays to compare the secretory activity associated with each virus^28^. SA11-G10-OSUv was associated with a larger percent increase in cross-sectional area than SA11-G10-OSUa (Figure 5D), suggesting that the wave inducing NSP4 from OSUv is associated with more fluid secretion than the non-wave inducing NSP4 from OSUa.

### NSP4 influences RV pathogenesis in mice

Our results in HIOs suggested that NSP4-OSUv and -OSUa differ in their ability to induce fluid secretion. Because fluid secretion is an integral aspect of RV pathogenesis, we sought to determine whether the disease phenotype associated with NSP4-OSUv and NSP4-OSUa would differ *in vivo*. We generated novel viruses using the recently developed D6/2 murine-like RV reverse genetics system^15^. We packaged viruses with either NSP4-OSUa (D6/2-G10-OSUa) or NSP4-OSUv (D6/2-G10-OSUv). Both strains grew to similar titers in MA104 cells; however, while the D6/2-G10-OSUv variant formed clear plaques in MA104 cells, D6/2-G10-OSUa failed to do so (Figure 6a).

**Figure 6:**
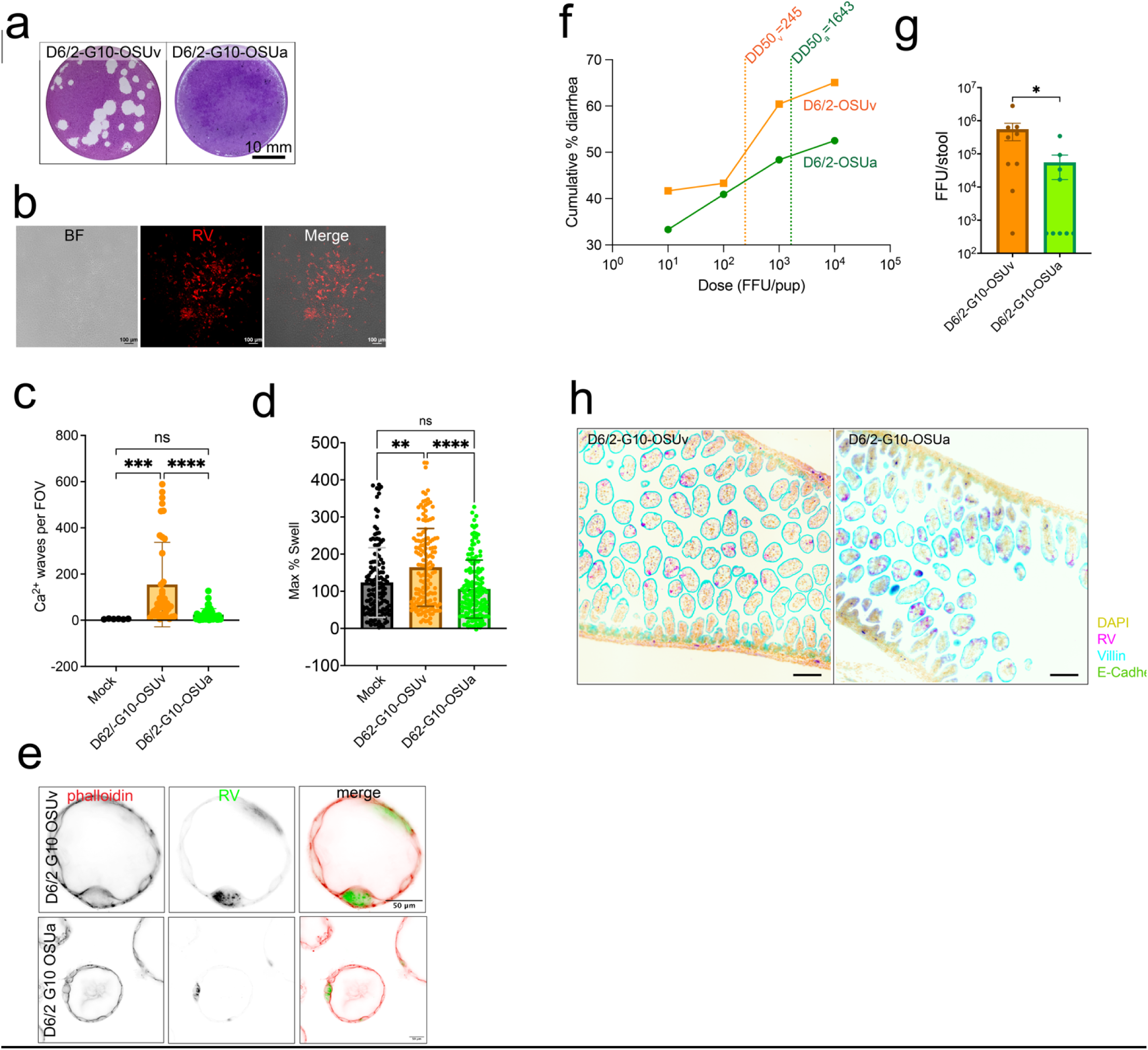
NSP4 is associated with Ca^2+^ waves, organoid swelling, disease, and shedding in a murine-like RV background. **a)** Representative images of plaque assays showing plaque clearance at 3 dpi in MA104 monolayers infected with D6/2-G10-OSUv, but non-lytic plaques in monolayers infected with D6/2-G10-OSUa. **b**) Representative brightfield and immunofluorescence image of monolayer infected with D6/2-G10-OSUa (red) showing the presence of infected foci despite the lack of plaque clearance. **c**) Quantitation of ICWs in MA104-GCaMP6s monolayers imaged every 1 min from 8-24 hpi. Data combined from 3 biological replicates. **d**) Max percent swell in mouse intestinal organoids (Balb/c jejunum) infected with the indicated RV strains. **e)** Detection of RV antigen (green) and filamentous actin (red) in mouse intestinal organoids. **f)** Litters of CD-1 mice were infected with the indicated dose of D6/2-G10-OSUv (*n*=40 pups across 4 litters) or -OSUa (*n*=63 pups across 4 litters) and followed for 5 days post-infection. The percentage of pups to develop diarrhea during the 5-day period was used to plot the cumulative incidence curves and estimate the dose required to produce disease in 50% of animals (diarrhea dose 50, DD50) for each virus. **g)** Infectious particles detected in stool from animals infected with 10^4^ FFU at 3 dpi by fluorescent-focus assay. n=9 pups per condition**. h)** Representative immunofluorescence images of RV (magenta), DAPI (yellow), villin (cyan), and E-cadherin (green) detected in intestinal epithelium from infected pups at 2 dpi. Scale bar = 100 um.

Nevertheless, detection of RV antigen by immunofluorescence showed that plaque-like foci still formed in monolayers infected with D6/2-G10-OSUa (Figure 6b). Thus, NSP4-OSUa in a D6/2 background impaired MA104 cell cytotoxicity in a manner we did not observe in the SA11 background. Like SA11 however, live Ca^2+^ imaging of infected MA104-GCaMP6s monolayers showed that OSUv-NSP4 was associated with a higher frequency of ICWs than OSUa-NSP4 in the D6/2 background (Figure 6c). To assess the secretory phenotype associated with the D6/2 strains, we first performed swelling assays in jejunal murine intestinal organoids (jMIOs). jMIOs infected with D6/2-G10-OSUv showed a larger percent increase in cross-sectional area than those infected with D6/2-G10-OSUa (Figure 6d), recapitulating our findings from SA11-infected HIOs. Immunofluorescence provided further evidence of successful infection (Figure 6e).

Next, we determined the diarrhea dose 50 (DD50) for both viruses in the CD1 (ICR) neonatal mouse model of RV diarrhea by infecting individual litters with 10^1^, 10^2^, 10^3^, or 10^4^ focus-forming units (FFUs) per pup. The estimated DD50 for D6/2-G10-OSUa was 1643 FFU, while it was just 245 FFU for D6/2-G10-OSUv (Figure 6f), representing a 6.7-fold decrease in the DD50 for the OSUv NSP4 variant. We also compared the quantity of infectious viral particles shed from litters infected with 10^5^ FFU/pup by performing focus-forming assays on stool samples collected at 3 days post-infection. Animals infected with D6/2-G10-OSUv shed, on average, 10-fold more virus than animals infected with D6/2-G10-OSUa (Figure 6g). Immunofluorescence staining of intestinal sections extracted from a subset of these animals provided further evidence of infection (Figure 6h).

## DISCUSSION

Through this work we established RV nonstructural protein 4 (NSP4) as the driver of RV-induced intercellular Ca^2+^ waves (ICWs). Further, these studies indicated that it is the NSP4 viroporin function, and production of intracellular Ca^2+^ signals, that correlates with ICWs. NSP4’s capacity to disrupt *intra*cellular Ca^2+^ is integral for ICW induction, suggesting ICWs occur not in response to canonical recognition of pathogen associated molecular patterns (PAMPs), but rather from aberrations in Ca^2+^ regulated signaling pathways. The targeted mutations between these pairs result in major differences in Ca^2+^ dysregulation, yet the pairs maintain 96-98% of their sequence homology. Despite the difference in the mode of activation, RNA sequencing suggests that the recognition of a pathogen through detection of aberrant signaling results in many of the same downstream transcriptional consequences. Mediators of innate immunity including CCL20, IL6, IL8, CSF1, NOD2, and TRAF1 are upregulated by the NSP4 variants capable of disrupting intracellular Ca^2+^ and causing ICWs (OSUv, SA11 NSP4-WT), but not in response to their counterparts that show diminished disruption of intracellular Ca^2+^ and fewer ICWs (OSUa, SA11 NSP4-ASDASA).

A local increase in cytosolic Ca^2+^ is a known instigator of ICWs^38,39^. However, how the elevation of cytosolic Ca^2+^ is sensed and translated into the release of purinergic signals remains an open question. Recent work established that intracellular Ca^2+^ levels modulate the activation of pannexin-1, a well-characterized ATP-permeable plasma membrane channel ^40^. Elevated intracellular Ca^2+^ activates CaMKII, which phosphorylates pannexin-1 and increases its ATP conductance. Alternatively, nucleotide release may occur not in direct response to elevated Ca^2+^, but as a result of disrupted Ca^2+^ regulated homeostatic processes. Ca^2+^ regulation of the actin cytoskeleton is one such process, conserved across biological kingdoms^41^. NSP4 Ca^2+^ mobilization causes actin remodeling^42^ which may serve as a mechanical stimulus, one of the longest studied ICW instigators^43,44^.

While NSP4-mediated intracellular Ca^2+^ dysregulation caused both transcriptional changes and ICWs, we have yet to determine the relationship between these two effects. Treatment with BPTU, a known inhibitor of RV ICWs^23^, did not ameliorate the transcriptional changes following NSP4 expression. This would suggest that these changes are not dependent on ICWs, but instead arise in direct response to NSP4-mediated intracellular Ca^2+^ dysregulation. However, in order to produce a level of NSP4 that mimicked an RV infection, we had to transduce cells with AAV-NSP4 at a high multiplicity. Despite the high multiplicity, the expression of the fluorescent reporter and immunoblot analysis indicate that only a subset of cells express an amount of NSP4 comparable to a RV infection. Nevertheless, nearly every cell is expressing *some* NSP4, thus likely experiencing some degree of viroporin-mediated Ca^2+^ disruption. In this setting, the transcriptional effects specific to ICWs may be masked by the NSP4-generated increase in Ca^2+^ signals. In a low multiplicity RV infection, there would be far more cells receptive to ICWs than expressing NSP4. We would predict the ICWs to have a proportionally greater effect in this setting. Furthermore, rotavirus nonstructural protein 1 (NSP1) antagonizes antiviral responses, which may offset some of the NSP4 induced transcriptional changes within infected cells^45,46^. This antagonism is not predicted to occur in uninfected neighboring cells, further augmenting the relative effect of ICWs.

Our studies characterized distinct Ca^2+^ signaling phenotypes associated with NSP4 from the virulent porcine RV strain, OSUv, and that from the attenuated porcine strain, OSUa. OSUv-NSP4 was also associated with a left-shifted diarrhea dose curve and more viral shedding *in vivo*, suggesting higher pathogenicity. It is possible that the propensity to cause diarrhea stems from accelerated growth kinetics. We have previously shown that the IP_3_R activation associated with ICWs facilitates viral spread *in vitro*^24^, and this may also contribute to the increased shedding associated with the ICW-prone OSUv-NSP4 observed here. However, OSUv-NSP4 was also associated with increased swelling in our HIO model, despite no detectable difference in viral yield. Because these experiments last just 12-18 hours and there is no trypsin in the culture media to activate viral progeny, viral spread would not be a contributing factor in HIO swelling. This suggests that ICWs may have pathogenic consequences both related to and independent of their ability to promote viral spread.

Through sequencing we found that OSUv-NSP4 and OSUa-NSP4 differ at just three out of 175 amino acids. Reversion of amino acid 172 from the threonine in OSUa-NSP4 to the isoleucine in OSUv-NSP4 was sufficient to recover the Ca^2+^ signaling phenotype associated with OSUv-NSP4. This includes both the P2Y1-independent intracellular Ca^2+^ signals, presumedly coming from NSP4 itself, as well as the P2Y1-dependent ICWs. Residue 172 is near the carboxy terminus in the cytoplasmic tail of NSP4. How this influences Ca^2+^ mobilization remains an outstanding question. Previous work has shown that this region of NSP4 binds both host proteins, including tubulin, and the viral protein VP6^47–49^. Since the differential Ca^2+^ phenotypes of NSP4-OSUv and -OSUa persist when NSP4 is expressed alone, they cannot be dependent on NSP4 interactions with other RV proteins. Thus, amino acid 172 must either influence Ca^2+^ signaling via an interaction with a host protein or by directly modulating Ca^2+^ conductance through the viroporin domain. Our initial hypothesis was that this residue was involved in a protein-protein interaction that influenced the rate of NSP4 exit from the ER into COPII vesicles and eventually the autophagosome-like compartment that co-localizes with RV viroplasms. In this way, the NSP4 cytoplasmic tail could indirectly reduce the NSP4 intracellular signals by regulating the amount of NSP4 in the ER with access to that organelle’s Ca^2+^ store. However, we detected no difference in the amount of NSP4 colocalized with general ER markers in cells infected with OSUv versus OSUa. Nevertheless, we cannot rule out that the NSP4 cytoplasmic tail sequence regulates a differential distribution to different ER subdomains, such as ER-exit sites, where its channel function may be reduced. While the structure of full-length NSP4 is not yet solved, it would offer insight into the role of the carboxy terminal domain in NSP4-mediated Ca^2+^ mobilization.

By infecting GCaMP6s-expressing human intestinal organoids with the virulent human RV strain, Ito, or the live-attenuated vaccine strain, RV1, we found a similar association between ICWs and virulence for human RV. The virulent strain induced a higher frequency of ICWs despite comparable, or even perhaps reduced, infectivity. While we have yet to attribute these differences to NSP4, these results suggest that ICWs are an important facet of human RV pathogenesis. In the work detailed here, we opted to focus on OSUv and OSUa NSP4 as the high sequence homology made for more efficient phenotype mapping. We predict that results from ongoing studies will imply a similar role for NSP4-induced Ca^2+^ signaling in human RV virulence.

Overall, this work identifies viroporin-mediated disruption of cytosolic Ca^2+^ as a novel signifier of pathogenic invasion and a key contributor to RV virulence. Viroporin activity induced transcriptional changes associated with canonical innate immunity, but also ICWs previously shown to promote viral spread^24^. ICW induction was associated with increased viral shedding and disease *in vivo*, suggesting that ICWs may confer a selective advantage to RV that, when combined with immune suppression by other viral proteins^17,45^, supersedes the deleterious effects predicted by innate immune activation. This implicates NSP4 and the pathways that respond to the associated Ca^2+^ aberrations as attractive therapeutic targets. Manipulation of NSP4 in live-attenuated vaccine strains may allow for attenuation without significant changes to structural proteins that constitute key RV antigens, promoting immunity without fulminant disease. Drugs that target the NSP4 viroporin itself, Ca^2+-^responsive proteins, or P2Y1-mediated ICWs may be useful antiviral and antidiarrheal therapeutics. Given that other enteric viruses, including caliciviruses and picornaviruses, encode Ca^2+^-conducting viroporins, these findings may have implications for human pathogens beyond RV^50,51^.

## Acknowledgements

This work was supported by National Institutes of Health Grants R01AI158683 and NIH R01DK115507 (PI: J. M. Hyser); S10OD028480 (PI: M. K. Estes) that supported purchase of the Zeiss LSM980-Airyscan 2 microscope; and P30DK56338 that supports the Texas Medical Center Digestive Diseases Center and the Gastrointestinal Experimental Model Systems (GEMS) core. NIH fellowship of authors are as follows: J.T.G was supported by NIH F30DK131828, the McNair Foundation M.D./Ph.D. Scholars Program and Histochemical Society Graduate Medical Trainee and Graduate Student Cornerstone Grant, F.J.S. was supported by NIH F31DK132942, K.A.E. was supported by NIH F32DK130288 and Histochemical Society Postdoctoral Keystone Grant. We thank Dr. Sashi Ramani for the RV1 rotavirus vaccine strain.

## METHODS

### Cell culture

MA104 (ATCC CRL-2378.1) and LLC-MK2 (ATCC CCL.7) cells were cultured in high-glucose Dulbeccos modified Eagle’s medium (DMEM) with 10% fetal bovine serum and 1X Antibiotic/Antimycotic (Invitrogen) at 37C with 5% CO_2_. Cells were passaged every 2-3 days.

### Rotaviruses

The parental OSUv and OSUa strains (gifts from Leonart Svensson), Ito strain (gift from Mary Estes), and RV1 strain (gift from Sashi Ramani) were propagated by inoculating MA104 cells at a low MOI (∼0.01) for 1 hr before rinsing and switching to DMEM with 1 ug/mL Worthington’s Trypsin for 3 days or until >90% of the monolayer showed cytopathic effect. Stocks then underwent 3 freeze-thaw cycles and aliquots were activated with 10 ug/mL Worthington’s Trypsin at 37C for 30 min before use. Virus stock titers were determined by plaque assay or fluorescence focus-forming assay on MA104 cells. All stocks were stored at -80C between uses. The recombinant SA11 and D6/2 strains were generated using established protocols for reverse genetics^15,52^.

### Human intestinal organoids

All HIOs used in these studies were acquired from the Gastrointestinal Experimental Modal Systems Core at the Texas Medical Center Digestive Diseases Center. HIOs were maintained in Wnt3a, R-spondin-3, Noggin, EGF (WRNE) culture media as previously described^23,28^. HIOs were differentiated for 3-5 days prior to RV infections as previously described^28,53^. For Ca^2+^ imaging, 3-dimensional HIOs were digested into a single-cell suspension by incubating in TrypLE (Gibco) for 5 min and vigorously pipetting before dispersing onto Collagen-IV-coated imaging bottom plates (Greiner)^54^. After 24 hours, monolayers were switched from WRNE to differentiation media. Monolayers were seeded 72 hours prior to RV infection or imaging. For rotavirus infection, rotavirus stocks were diluted in CMGF- and a single score was introduced along the diameter of the monolayer using a 23-gauge needle. Monolayers were inoculated for 2 hrs, rinsed, then incubated in differentiation media.

### Live calcium imaging

MA104, LLC-MK2, and HIO lines stably expressing the cytosolic calcium indicator GCaMP6s were generated using lentivirus transduction as previously described^36,53,54^. Monolayers were seeded in 8-well, imaging-bottom chamber slides (Ibidi) and allowed to grow for 48 hours or until confluent. A single scratch was introduced down the length of HIO monolayers immediately prior to infection. Monolayers were mock- or RV-infected at a low multiplicity of infection (MOI 0.005-0.05) for 1 hr. For HIO studies, mock inoculum included mock-infected, trypsinized MA104 lysate equalized to the volume of virus stock included in RV infections. The inoculum was removed, monolayers rinsed, and media switched to phenol-free imaging medium (FluoroBrite, Gibco). 8 fields-of-view (FOV) were selected at random for each well. Using a 20X Plan Apo objective, 50 ms exposures at 50% light source power were acquired on the FITC line every 1 min, and TRITC line every 10 min, from 7-25 hpi. Slides were maintained in a stage-top, temperature- and humidity-controlled chamber with 5% CO_2_. For experiments involving RV strains that did not express a fluorescent reporter, following live imaging, monolayers were fixed with 4% formaldehyde and RV-infected cells were identified via immunofluorescence as described below.

### Image analysis

For the analysis of image series from long-term calcium imaging experiments we used the open-source platform *FIJI*^55^. FITC (GCaMP6s) and TRITC (RV mRuby) acquisitions were split into separate stacks and processed individually. Each image in the FITC time series was subtracted from the next to determine the change in fluorescence at each pixel between acquisitions (ΔF). A minimum ΔF threshold was applied (+600 relative fluorescent units) before detecting features based on a minimum size (10^4^ contiguous microns), which represent ICWs. The total number of ICWs within each field-of-view were compared across treatment conditions.

Antibodies, stains, dyes (supplemental table 1)

| Type | Target | Species | Supplier | Cat. # / ID | Dilution |
| --- | --- | --- | --- | --- | --- |
| Primary | SA11 rotavirus NSP4 aa114-135 | Guinea Pig | AB Clonal | Custom | 1:1000 |
| Primary | SA11 rotavirus NSP4 | Mouse | Gift from Dr. Mary Estes | MAb B4/2, purified | 1:500 |
| Primary | SA11 rotavirus | Rabbit | Gift from Dr. Mary Estes | Rabbit pool, exp 42 | 1:5000 |
| Primary | SA11 rotavirus vNSP2 | Mouse | Gift from Dr. Mary Estes |  | 1:2000 |
| Primary | Phospho-Ezrin | Rabbit | Cell Signaling Technologies | 3726 | 1:500 |
| Primary | E-Cadherin | Rabbit | GeneTex | 1004433 | 1:200 |
| Primary | Sec61β | Rabbit | Invitrogen | PA3-015 | 1:500 |
| Primary | GAPDH | Mouse | Novus | NB-600-502 | 1:5000 |
| Primary, conjugated | Villin | Mouse | Santa Cruz | Sc-58897 | 1:500 |
| Secondary | Anti-rabbit Alexa Fluor 488 | Donkey | Rockland | 611-741-127 | 1:1000 |
| Secondary | Anti-rabbit Alexa Fluor 555 | Donkey | Rockland | 606-142-129 | 1:1000 |
| Phalloidin 555 | Actin | *Amanita phalloides* | ThermoFisher | A34055 | 1:1000 |
| DAPI | dsDNA |  | Invitrogen | R37606 | 2 drops per mL |

### Immunofluorescence

Culture media was removed from MA104 or HIO monolayers before rinsing with 1X PBS. Monolayers were fixed for 30 min with 4% formaldehyde then washed 3 times with 1X PBS. HIO monolayers were incubated in 50 mM NH_4_Cl for 30 min following fixation. Monolayers were blocked for 1 hr at room temperature or overnight at 4C with 5% bovine serine albumin in 1X PBS, then permeabilized for 30 min with 0.01% Triton X-100 in 1X PBS. Monolayers were then incubated in primary antibodies diluted to the indicated concentrations in 1X PBS overnight at 4C, washed 4 times with 1X PBS, and incubated with the indicated fluorescently conjugated secondary antibodies for 1 hr at room temperature. Monolayers were again washed 4 times with PBS and maintained at 4C in 1X PBS until imaged.

### Western blot

Confluent mock-infected, RV-infected, or AAV-transduced MA104 cell monolayers seeded in 24-well culture plates were incubated in 150 uL RIPA buffer (50 mM Tris, 150 mM NaCl, 1% Nonidet P-40, 0.5% sodium deoxycholate, 0.1% sodium dodecylsulfate, 1 EDTA-free protease inhibitor tablet (Sigma-Aldrich), pH 7.4) and subjected to one freeze-thaw cycle. Where indicated, a 30 uL aliquot of lysate was transferred to a clean 1.5 mL microcentrifuge tube and incubated with 1 uL Endo-H (NEB) for 45 min in a 37C bead bath. 20% volume of 5X SDS-PAGE sample buffer was added to each lysate before incubating at 100C for 10 min. 30 uL of lysate was loaded per well of a pre-cast 4-20% polyacrylamide gel. Gels were loaded into an electrophoresis rig and run at 120V for 75 min. Protein was transferred to a nitrocellulose membrane using a semi-dry transfer apparatus. Membranes were allowed to dry completely, re-hydrated for 2 min with 1X PBS, and blocked for 1 hr using Intercept Blocking Buffer (Licor). Primary antibodies were diluted in 1X PBS with 0.2% Tween-20 and 10% Intercept Blocking Buffer by volume and incubated overnight. Membranes were washed 4 times with 1X PBS 0.2% Tween-20 before incubating secondary antibodies diluted 1:20,000 in 1X PBS for 1 hr. Membranes were rinsed an additional 4 times with 1X PBS 0.2% Tween-20, once with 1X PBS, then dried and imaged on a Licor Odyssey CLx scanner.

### RV growth curves

Confluent MA104 cell monolayers were infected with the indicated strains of RV for 1 hr at MOI 0.5. The inoculum was removed, and monolayers were washed with 1X PBS before replacing with serum-free high-glucose DMEM. At 1, 3, 6, 9, 12, and 24 hpi, monolayers underwent 3 freeze-thaw cycles, were transferred to 1.5 mL microcentrifuge tubes, centrifuged to remove cell debris, and incubated for 45 min at 37C with 10 ug/mL Worthington trypsin. RV yield was assessed by plaque assay as described below.

### Adeno-associated viruses

Adeno-associated viral vectors were synthesized and packaged commercially (VectorBuilder). For imaging experiments, 5x10^4^ MA104-GCaMP6s cells were seeded on 8-well imaging bottom chamber slides (Ibidi). 12 hours post-seeding, monolayers were transduced with 50,000 genome copies AAV per cell diluted in high-glucose DMEM with 10% fetal bovine serum. Transduction medium was left on monolayers for 24 hours, rinsed once with 1X PBS, then switched to phenol-free cell culture medium (FluoroBrite+, Gibco). Live Ca^2+^ imaging was performed from 32-50 hrs post-transduction as described above. For Western blots, 24-well cell culture plates were seeded with 1.5x10^6^ MA104 cells per well (Corning) and transduced 12 hrs post-seeding. Lysates were harvested with 150 uL RIPA buffer and processed as described above.

### Swelling assays

HIOs were switched to differentiation media 2 days post-passage. After 3-5 days of differentiation, HIOs were removed from basement membrane matrix using 500 uL ice-cold 1X PBS per well and spun for 4 min at 4C. Supernatant was removed and pellets resuspended in 1 mL 1X PBS before a second spin. Pellets were then resuspended in 200 uL inoculum consisting of 5x10^6^ PFU of trypsin-activated RV or tryspinized uninfected MA104 cell lysate diluted in CMGF^-^ for 1 hr at 37C. HIOs were pelleted, washed once with PBS, resuspended in 25% basement membrane matrix diluted in phenol-free differentiation media, and plated on an imaging bottom 8-well chamber slide (LabTek) pre-coated with a thin layer (∼4 uL) of basement membrane matrix. The slide was then loaded into a humidified stage-top incubation chamber maintained at 37C with 5% CO_2_ on a Nikon TiE inverted epifluorescent microscope. Nikon NIS-Elements software was used to acquire bright-field images of 50+ HIOs per condition every 5 minutes for 18 hrs, and 555 nm epifluorescent images every 20 min for experiments using mRuby-tagged RV strains. FIJI was used to measure the initial and maximal cross-sectional area of each HIO, and to calculate the percent change (maximum percent swell). MIOs were infected in suspension for 2 hrs 1-day post-passage using the same method.

### Yield assay in HIOs

Differentiated HIOs were removed from basement membrane matrix using ice-cold PBS, pelleted, washed in 1X PBS, and resuspended in 200 uL RV or mock inoculum as described above. After 2 hrs, HIOs were pelleted, washed, and half were harvested for the 2 hr titer, half resuspended in 30 uL basement membrane matrix until the 24 hr timepoint. To determine the viral yield, HIOs and supernatant underwent 3 freeze-thaw cycles before trypsin-activation and titration by plaque assay.

### Plaque assay

RV infected lysates (cells and supernatant) underwent 3 freeze-thaw cycles and activation with 10 μg/mL Worthington Trypsin for 30 min at 37C before 10-fold serial dilution. MA104 cells were grown to confluency in 6-well plates and switched to serum-free medium for 24 hrs pre-infection. Wells were infected with 200 μL of each serial dilution in duplicate for 1 hr at 37C with gentle shaking every 15 min to ensure even distribution. Monolayers were rinsed once with 1X PBS and media replaced with a semi-solid overlay consisting of 1.2% Avicel in serum-free DMEM supplemented with DEAE dextran and 1μg/mL Worthington’s Trypsin. After 72 hrs, overlay was removed, and monolayers stained with crystal violet.

### Fluorescence focus assay

A 96-well cell culture plate was seeded with 1.5x10^4^ MA104 cells per well and grown to confluency. 24 hours prior to infection, plates were switched to serum-free high-glucose DMEM. RV stocks were activated with 10 μg/mL Worthington Trypsin for 30 min at 37C before 4-fold serial dilution in serum-free high-glucose DMEM. Each well was inoculated with 100 μL for 1 hr at 37C. The inoculum was removed, wells rinsed with 1X PBS, and media replaced. At 18 hpi, the media was removed and wells rinsed with 1X PBS before fixation with 100% methanol for 20 min at 4C. Wells were rinsed 3 times with 1X PBS, incubated overnight with 50 uL of primary antibody solution, rinsed another 3 times, then incubated for 1 hr at room temperature with secondary antibody solution. The secondary antibody solution was removed, and nuclei stained with NucBlue Fixed Cell Stain per the manufacturer protocol. Monolayers were washed and maintained in 100 uL 1X PBS per well. A 10X/0.30 Plan Fluor objective on a Nikon Eclipse Ti epifluorescent microscope was used to acquire a 7.5 x 7.5 mm stitch image covering the entirety of each well. Rolling ball background subtraction and thresholding was performed in FIJI before quantifying the number of infected cells per well via the particle analysis plug-in.

### RNA Sequencing

1.5x10^5^ MA104-GCaMP6s cells were seeded per well in a 24-well cell culture plate. 12 hrs post-seeding, adeno-associated virus encoding NSP4 and a fluorescent reporter was added to indicated wells (5x10^4^ genome copies per cell) as described above. At 72 hrs post-transduction, the plates were imaged to verify expression of the fluorescent reporter before column-based isolation of RNA (Qiagen, RNeasy Mini Kit) with on-column DNA digestion. RNA quality and quantity was assessed by fluorimetry (Qubit Extended Range Assay Kit, ThermoFisher) and stored at -80C. Isolation of mRNA was achieved via poly(A) enrichment before library preparation (Ultra II Directional RNA library prep kit, New England Biolabs). Libraries were sequenced on an Illumina NovaSeq with 40M 150bp paired end reads per sample.

Read quality was assessed using FastQC v0.12.1^56^ and MultiQC v1.19^57^. Reads were aligned to the Chlorocebus sabaeus genome assembly 1.1 accessed via *Ensembl* (GCA_000409795.2)^58^ using STAR aligner v2.7.11a^59^. Expression of the AAV transgene was estimated by quantitating mScarlet transcripts using the *kallisto* v0.46.1 pseudoaligner^60^. Read counts were compared using PyDESeq2^61^ v0.4.4 and GSEApy^62^ v1.1.1.

### Mouse intestinal organoids (MIOs)

All MIOs used in these studies were acquired from the Gastrointestinal Experimental Modal Systems Core at the Texas Medical Center Digestive Diseases Center. MIOs were maintained in Advanced DMEM/F12 (Gibco) supplemented with 1X GlutaMax (Gibco), 0.01M HEPES, 100 U/ml penicillin-streptomycin (Gibco), 20% Rspo-1 conditioned medium, 5% Noggin conditioned medium, 1mM N2 supplement (Invitrogen), B27 supplement (Invitrogen), 1mM N-acetylcysteine (Sigma-Aldrich), and 50 ng/mL recombinant murine epidermal growth factor (Invitrogen). Every 3-4 days, MIOs were passaged by removing maintenance media, adding 300 uL 0.05% trypsin, and mechanically breaking up basement membrane matrix by gently pipetting up and down before incubating for 5 min at 37C. 500uL of CMGF- was then added and the suspension was vigorously pipetted 20x to promote crypt fission. The suspension was pelleted, washed once with 1X PBS, then resuspended in basement membrane matrix before replating.

### Diarrhea dose 50 in RV mouse model

All experiments were approved by the Institutional Animal Care and Use Committee at the Baylor College of Medicine (protocol AN-6903) and performed in accordance with the National Institutes of Health Guide for the Care and Use of Laboratory Animals. For all mouse experiments, litters including female and male animals in naturally occurring ratios were used. CD-1 dams with litters of males and females in natural ratios were purchased from the Center for Comparative Medicine at Baylor College of Medicine. Pups were housed with their dam in standard cages in a BSL2 facility with food diet (7922 NIH-07 Mouse diet, Harlan Laboratories) and water provided *ad libitum* throughout the experiment. Individual litters were randomly assigned for inoculation with 10, 10^2^, 10^3^, or 10^4^ FFU/pup of D6/2 G10-OSUv or D6/2 G10-OSUa at postnatal day 3. Pups were infected by oral gavage with inoculum consisting of virus stock diluted with DMEM to a final volume of 50 uL per animal. Filter-sterilized green food-grade dye was included in the inoculum to visualize delivery to the stomach.

Litters were monitored for 5 days post infection with weights and stool assessed daily. At the time of assessment, pups were gently palpated on the abdomen to encourage defecation. Stool was scored using a 4-point scale for consistency, color, and quantity as previously described^19^. Stool was collected and stored at -80C for later assessment of viral titer, as well as secondary scoring by a blinded observer. Pups with weights that remained >2 standard deviations below the mean weight of the litter were excluded from analysis. The amount of virus required to induce disease in 50% of pups (diarrhea dose 50, DD50) was estimated using methods previously described^63^.

### Assessment of viral shedding

Stool from animals infected with 10^4^ FFU was dissociated by adding 40 uL of 1X PBS with Ca^2+^ and Mg^2+^ after thawing. Each sample was then sonicated for 30 min, diluted 1:100 with serum-free DMEM, and activated with 10 ug/mL Worthington Trypsin for 30 min at 37C. After activation, samples were serially diluted 1:4 with serum-free DMEM. Fluorescence focus assays were performed as described above.

### Mouse intestine histology

CD-1 litters with males and females in natural ratios were inoculated with 10^5^ FFU/pup of D6/2 G10-OSUv or D6/2 G10-OSUa at postnatal day 3. At days 3, 4, and 5 post-infection, 3 animals were euthanized from each litter for histological examination. Following euthanasia, the small intestine was extracted, flushed with 1 mL 1X PBS delivered via 23G needle inserted into the proximal end of the duodenum, then flushed with 10% formalin by the same method. The intestine was then rolled from the proximal to distal end and placed in a tissue cassette. The tissue cassettes were then immersed in 10% formalin overnight before switching to 70% ethanol. The samples were paraffin embedded and sectioned onto glass slides.

One slide from each animal underwent hematoxylin/eosin staining for general histopathological examination and one was used for detection of RV by immunofluorescence. Slides used for immunofluorescence were deparaffinized and rehydrated in xylene followed by gradient ethanol and distilled water. Slides were incubated in sodium citrate buffer (10 mM sodium citrate, 0.05% Tween 20, pH 6.0) for 15 min in a high-pressure pressure cooker, followed by 15 min of depressurization and cooling. The slides and buffer were then incubated on ice for 30 min then washed in 1X PBS for 5 min. The tissue was blocked for 1 hr at room temperature with 10% normal goat serum in 1X PBS before a 24 hr, 4C incubation with primary antibodies diluted in 1X PBS. Primary antibody solution was removed, slides rinsed with 1X PBS 3 times for 3 min, then secondary antibodies added for 1 hr at room temperature. Secondary antibody solution was removed and replaced with NucBlue Fixed Cell Stain for 10 min at room temperature per manufacturer protocol. After 3 additional 3 min washes with 1X PBS, excess liquid was allowed to evaporate before applying mounting media (ProLong Gold) and coverslips. Stained tissue sections were imaged using a Zeiss AxioScan Z1 whole slide scanner with a 20X/0.80 objective.

---

**Supplemental Figure 1:**
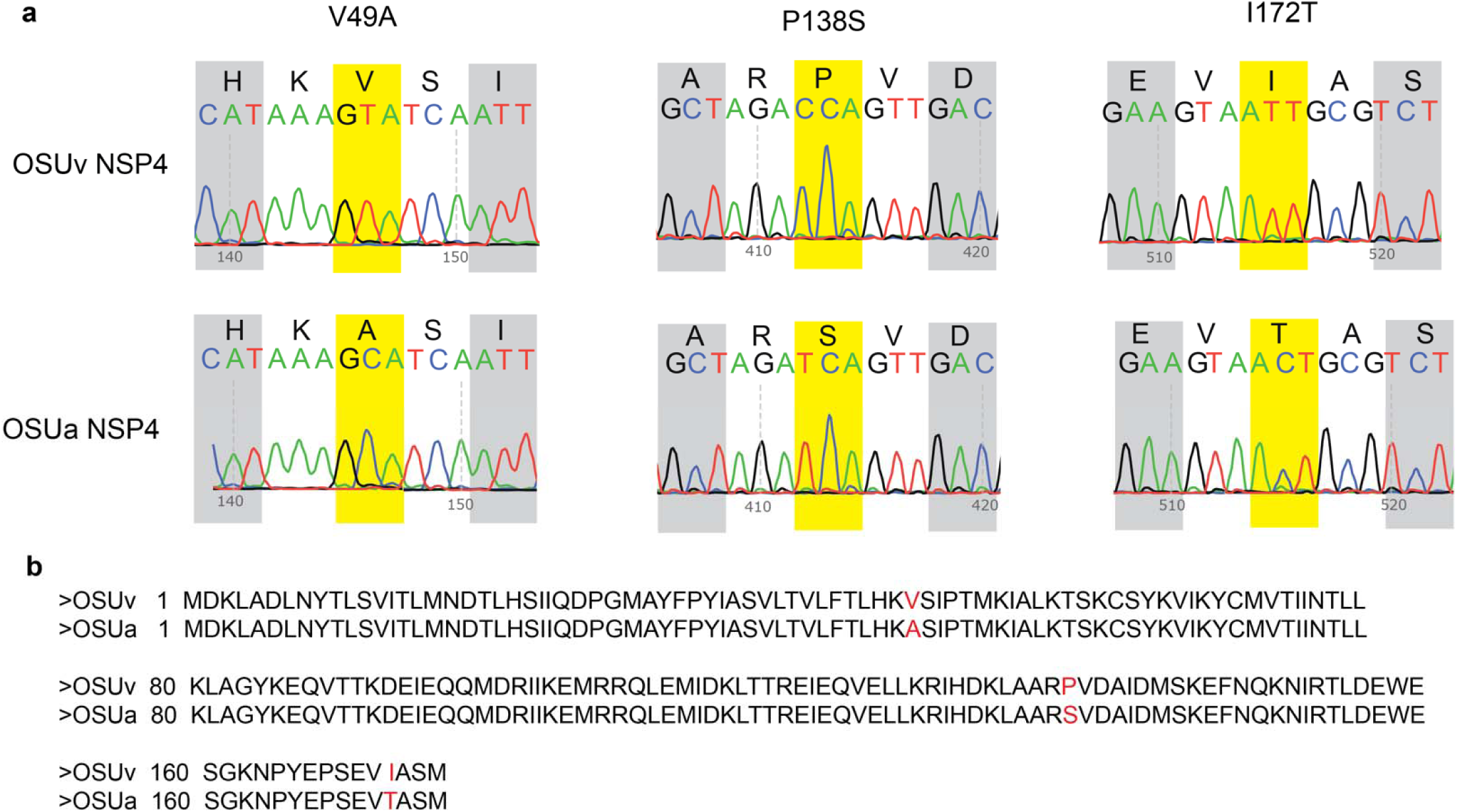
NSP4 from OSUv and OSUa differ by 3 amino acids. **a.** dsRNA from OSUv and OSUa rotavirus was isolated, reverse transcribed, amplified, and column-purified for Sanger sequencing, which revealed 3 polymorphisms in OSUa (yellow). **b.** Sequence alignment of full-length NSP4 from OSUv and OSUa showing the 3 resultant amino acid polymorphisms (red).

**Supplemental Figure 2:**
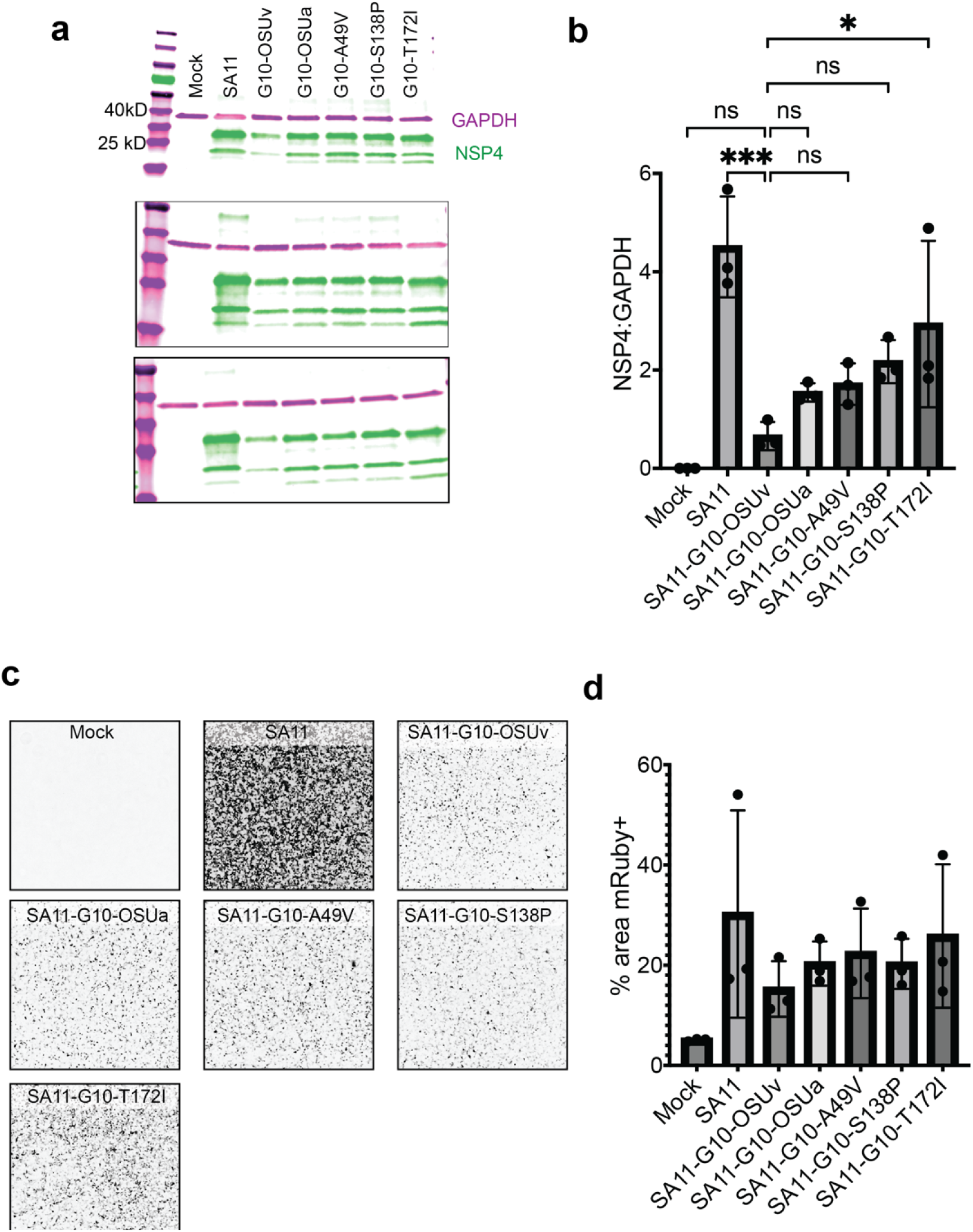
Comparison of NSP4 and fluorophore expression from recombinant RV strains. **a.** Western blot for GAPDH (magenta) and NSP4 (green) detected in lysates from MA104 cells infected with the indicated strains of RV at MOI 1 at 12 hpi. Three biological replicates were run independently (vertical panels). **b**. The intensity ratio of NSP4 to GAPDH was compared between viruses. Each point represents the ratio for the indicated sample from a single biological replicate. Data was analyzed by one-way ANOVA followed by Dunnett’s multiple comparisons tests between SA11-G10-OSUv and all other samples. **c**. Representative fluorescent images of monolayers infected with the indicated RV strains. Prior to lysis (11.5 hpi), monolayers were imaged to compare the expression of mRuby, encoded downstream of NSP3 on gene 7 for all RV strains included. Scale bar = 1 mm. **d.** Comparison of the percentage of each monolayer showing mRuby positivity. 3 images were collected per group (one per biological replicate). mRuby positivity was quantified following flat-field correction, background subtraction, and uniform thresholding.

**Supplemental Figure 3:**
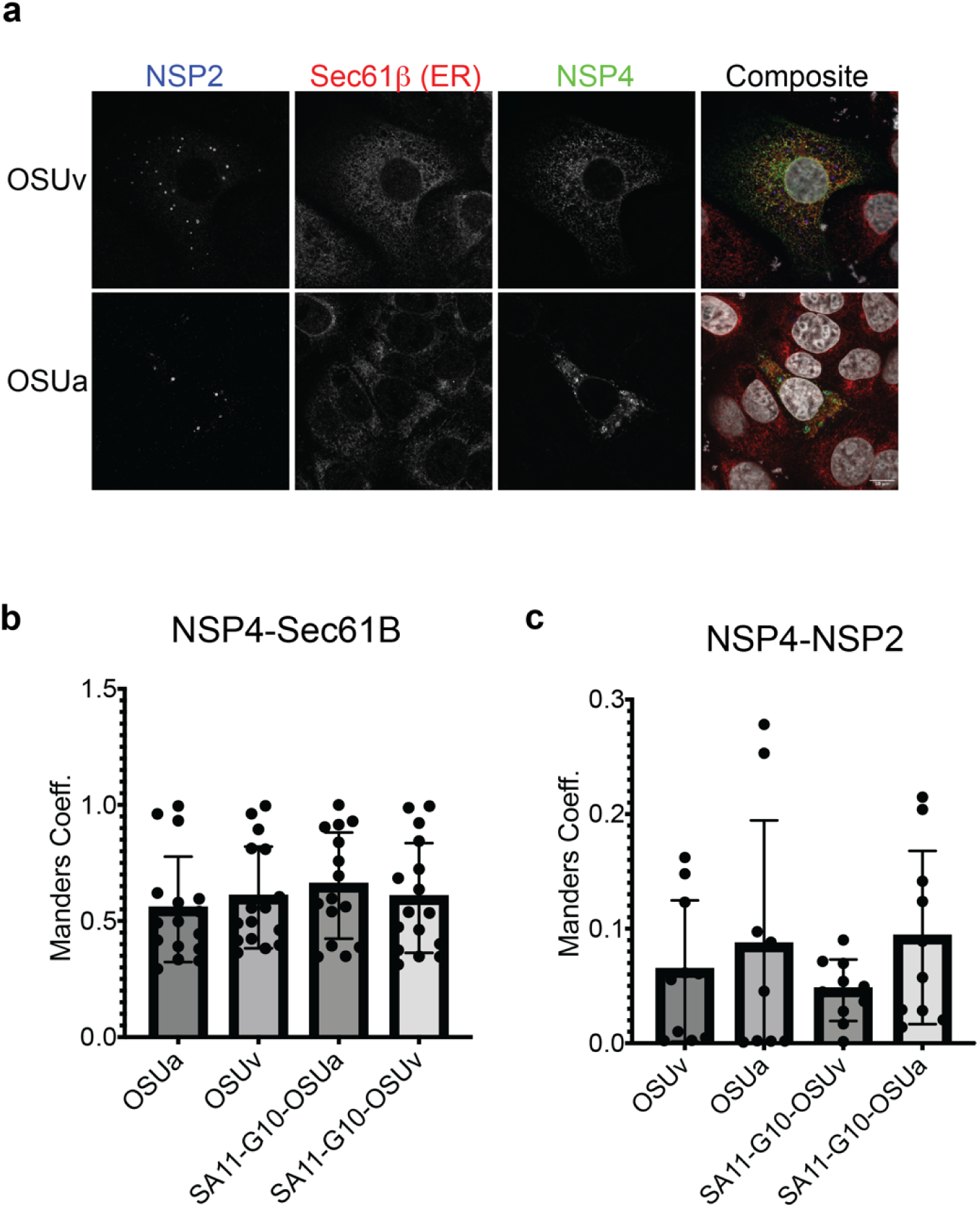
Assessment of NSP4 colocalization with viroplasms (NSP2) or ER (Sec61β) by immunofluorescence. **a.** Representative images of MA104 cells infected with the indicated viruses at MOI 1 before fixing at 12 hpi and detecting the indicated proteins by immunofluorescence. 63X magnification. **b.** Manders coefficient estimating colocalization between OSUa and OSUv NSP4 with Sec61β by immunofluorescence. n=16 cells per condition, 3 biological replicates **c.** Manders coefficient estimating colocalization between OSUv and OSUa NSP4 with NSP2, a marker of viroplasms. n=9 cells per group, 3 biological replicates. Shapiro-Wilk test for normality (p<0.05 for 3/4) followed by Kruskal-Wallis (p>0.05).

**Supplemental Figure 4:**
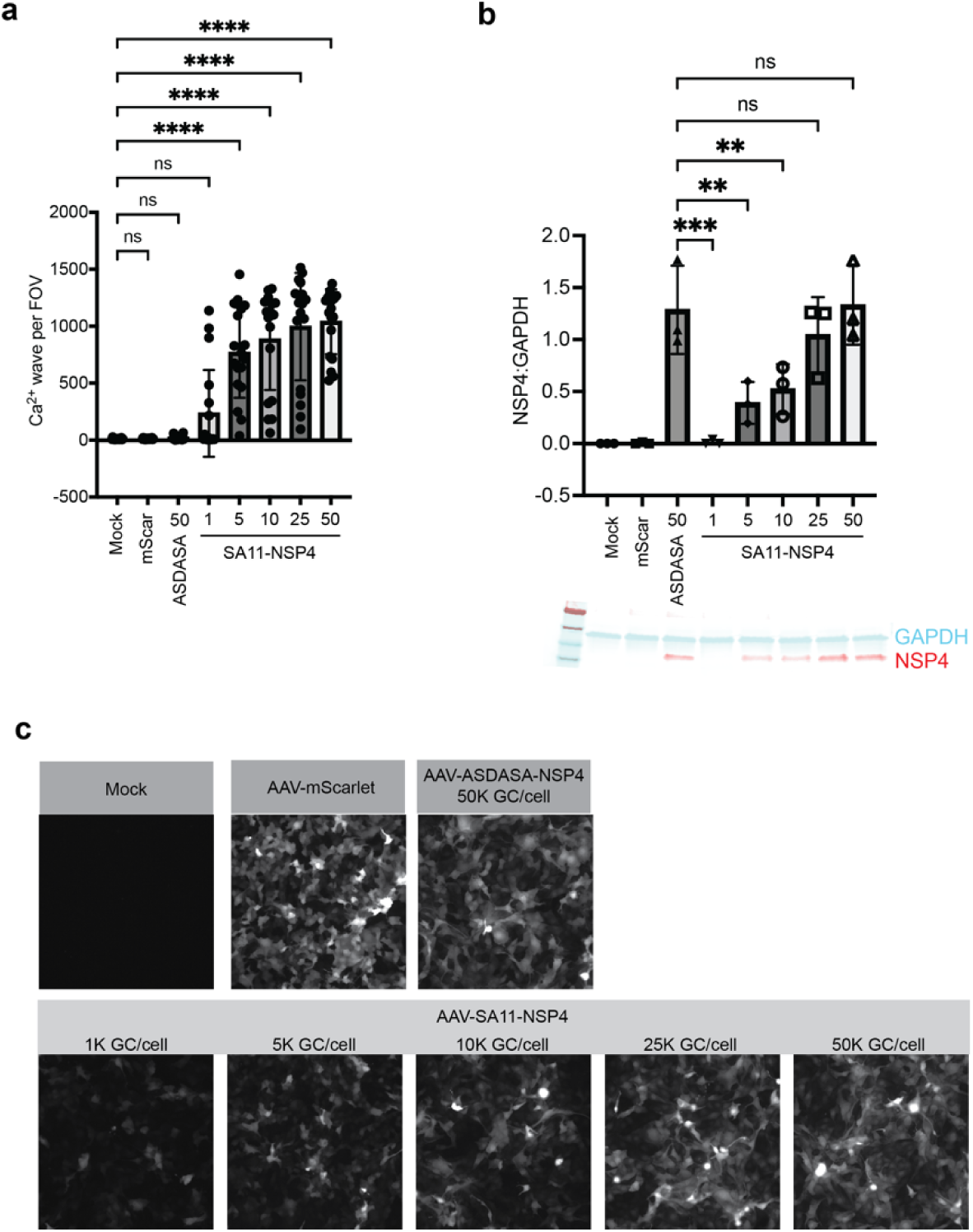
Comparison of SA11 NSP4 and ASDASA NSP4 using descending multiplicities of AAV. **a**. ICWs detected in monolayers of MA104-GCaMP6s cells transduced with the indicated AAV constructs at a multiplicity of 1k, 5k, 10k, 25k, of 50k genome copies per cell as indicated. Monolayers were imaged once per minute at 8 locations per condition from 32-50hpt. Results were combined from 3 independent experiments. **b**. Comparison of NSP4 level normalized to GAPDH for each condition. Below the graph is an image of a representative nitrocellulose membrane from one of three biological replicates with the lanes corresponding to the labels on the graph above. Indications of statistical significance represent results from Dunnett’s multiple comparisons following Kruskal-Wallis test.

